# Efficient exploration of sequence space enables rapid generation of functional genome editors

**DOI:** 10.64898/2026.08.16.745112

**Authors:** Nicholas W. Hughes, Sourab Kulkarni, Grant Goldman, Julia Marsiglia, Sunit Jain, Kaitlyn Spees, Becky Xu Hua Fu, Kia Vaalavirta, Muneaki Nakamura

## Abstract

The problem of how protein sequences translate into defined functions remains largely unsolved despite decades of progress. New methods to efficiently explore protein sequence space will help to shed light on these sequence-function relationships, particularly for complex protein function. Here, we describe an approach to create novel, functional proteins through the integration of deep mutational scanning, structural analysis, and evolutionary mining within prompts for a generative protein language model (PLM). We demonstrate the utility of this approach with the generation of novel compact RNA-guided nucleases. This approach is highly efficient, resulting in active nucleases with ∼40% sequence divergence relative to natural proteins and activity equivalent to or exceeding by up to ∼3X that of other compact nucleases at multiple endogenous loci in human cells. The approach described here is rapidly deployable and produces new sequences that will serve as scaffolds for further exploration of complex protein functionality, as well as substrates for novel genome engineering applications.

## INTRODUCTION

Due to the extreme diversity of possible protein sequences, unguided experimental exploration is intractable for *de novo* protein design, necessitating computational approaches to constrain assessed sequence space. Over the past few decades, significant progress has been made in producing functional proteins with defined structures, and more recently, with designed binding affinities^1^. However, translating these approaches for more complicated enzymatic functions remains non-trivial.

Recent work has turned to the use of language models (LMs), which seek to directly infer sequence-function relationships, bypassing the need for an explicit structural intermediate. This has resulted in the template-based creation of proteins encompassing a range of more complex functions. For example, a central application of language models has been on the design and generation of new RNA-guided nucleases^2–4^, which are widely applicable across research and biotechnological applications. Notably, this enzymatic class is particularly challenging to design since it consists of multi-domain proteins that must recognize and bind multiple nucleic acid substrates across distinct conformational states.

While these proof-of-concept reports highlight the potential of LM-based approaches for generating novel complex proteins, they often involve proprietary models and bespoke finetuning, which produce sequences that bear only a modest (∼20%) sequence divergence from the input protein templates. Exploring more diverse regions of sequence space will allow more comprehensive probing of sequence-function relationships, shedding new light on protein function outside of naturally occurring sequences or closely derived variants identified through existing LM-based approaches.

Here we describe a framework for prompt design for protein language model (PLM)-driven novel protein generation. Based on deep mutational scanning^5^ (DMS) data collected in human cells, we created a multi-modal functional, structural, and evolutionary prompt generation approach that included steering algorithms^6^ to bias generation toward more diverse and high-activity regions of sequence space. This approach yields synthetic <u>**A**</u>crobat <u>**c**</u>ompact <u>**e**</u>ditors (ACEs), which can robustly edit the human genome while demonstrating high sequence divergence, retaining only 57-61% identity to the closest naturally occurring proteins.

## RESULTS

### Generative Design Approach

We reasoned that combining information across complementary layers would enable us to identify a core set of residues to prompt a generative PLM, which would potentially enable greater diversity of generated sequences while maintaining high levels of activity (**Fig. 1a**). Specifically, we sought to incorporate functional, structural, and evolutionary information into a single prompt to use as input for ESM3^7^, which is a generative protein language model used previously to engineer novel variants of GFP. As a test case of this approach, we turned to the generation of novel nucleases using the TnpB^8,9^ family as a template. At roughly 400 amino acids it is less than a third of the size of SpCas9 and leaves substantial cargo capacity within a single adeno-associated viral vector. TnpB is under active development as a genome editing tool in human cells^10–12^, and its mechanism has been resolved structurally in both guide-bound and target-bound states^13,14^. TnpB is therefore a therapeutically motivated enzyme for which preliminary data already exist.

**Figure 1.**
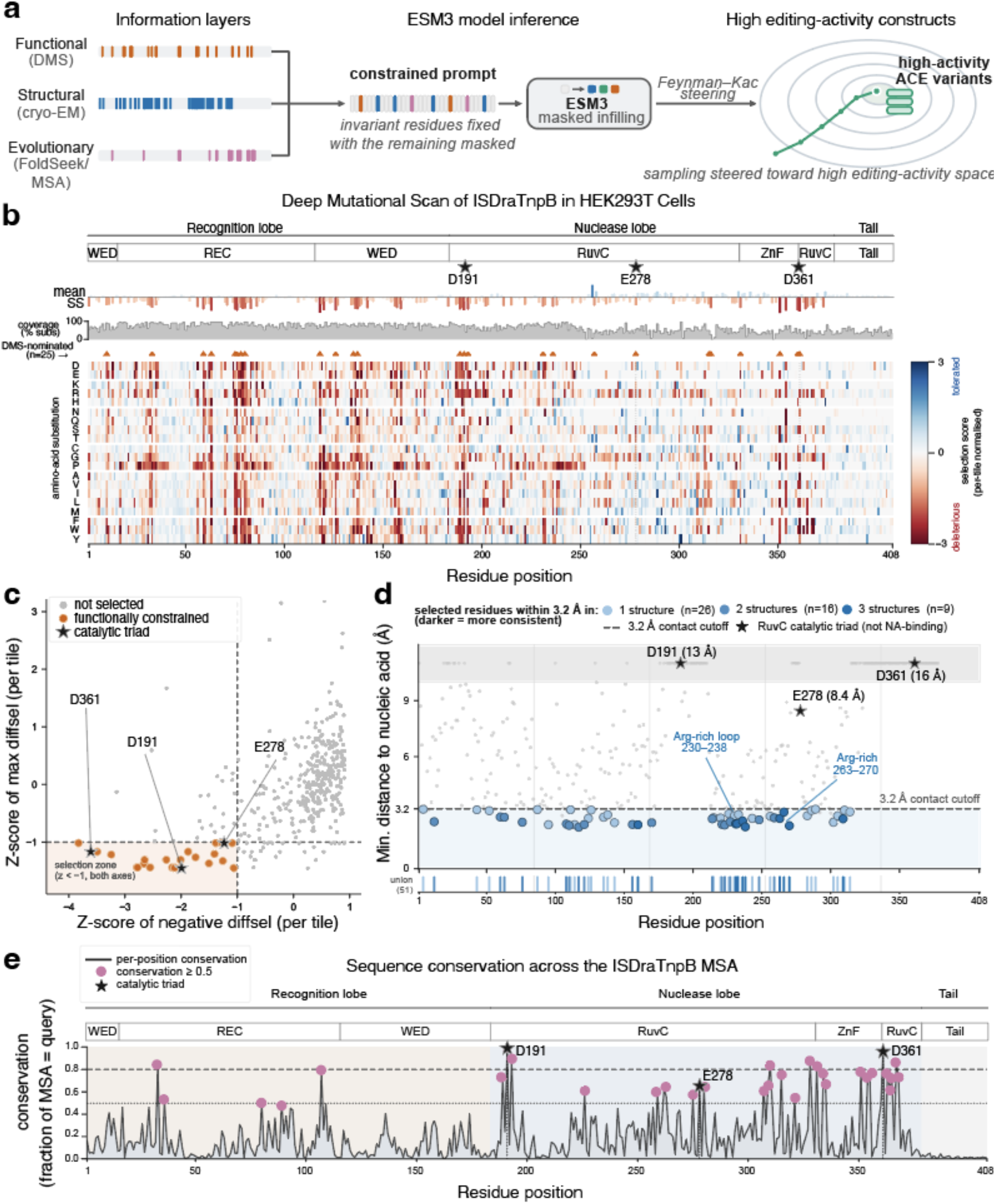
Generative design using biologically informed prompt constraints. **(a)** Three evidence layers - functional (DMS, orange), structural (cryo-EM nucleic acid contacts, blue) and evolutionary (Foldseek, purple) - nominate 151 invariant residues that define a constrained ESM3 prompt. Masked positions are infilled via either an unsteered or Feynman-Kac (FK) steering approach, yielding ACE designs. **(b)** Deep mutational scan of DraTnpB in HEK293T cells. Per-mutation selection scores (red deleterious, blue beneficial) for each possible single variant are depicted. **(c)** Functional selection in which residues depleted (z < -1) on both negative and maximum differential selection metrics across at least 2 experimental batches. These residues are then expanded to constrain all residues within 4 Å. **(d)** Structural constraint in which the minimum distance to nucleic acid across three cryo-EM structures (8EXA,8H1J,8BF8) was computed for each residue. Residues within 3.2 Å were constrained. **(e)** Evolutionary constraint in which the per-position MSA conservation score was computed after querying FoldSeek for structurally similar proteins. All residues that were conserved in >= 50 % of variants within the MSA were constrained.

We built upon this preliminary data by performing a deep mutational scan of the ISDra2 TnpB from *D. radiodurans* (DraTnpB) directly in human cells. We constructed a lentiviral GFP-ON reporter, containing an out-of-frame EGFP cassette with an upstream DraTnpB target site, such that 3n+2 indels generated by error-prone repair restore the EGFP reading frame and activate fluorescence (**Supplementary Fig. 1a**). We delivered a pooled library encoding all 7,752 single amino acid substitutions of DraTnpB into a HEK293T line carrying this reporter and the cognate guide RNA (gRNA) targeting the GFP-ON sequence. We separated edited from unedited cells by FACS, and quantified variant abundance in each fraction by tiled amplicon sequencing to estimate the fitness effect of each variant **(Fig. 1b, Supplementary Fig. 1b**,**c)**. Twenty-five residues, including the RuvC catalytic triad (D191, E278 and D361) were identified as critical for TnpB function with a stringent selection window (**Fig. 1c**). We hypothesized that the function of these residues may be dependent on their local geometry within the protein structure and therefore fixed residues in the ligand-bound structure (PDB 8EXA) within 4 Å of each of these critical residues, yielding 61 functionally constrained positions within the functional layer.

We further leveraged structural information for the subsequent structural layer by fixing residues that engage the gRNA and target DNA. Across three cryo-EM structures of DraTnpB spanning duplex and target-bound states, we identified 51 residues lying within 3.2 Å of a nucleic acid in at least one structure (**Fig. 1d**). Notably, this layer formed a distinct set of residues compared to the functional layer, with the catalytic triad absent from this set, while the arginine-rich loops lining the gRNA interface (residues 230 to 238 and 263 to 270) were captured, yet scored as individually substitutable in the mutational scan. Overall, this indicates that the functional and structural layers report on different, complementary determinants of protein fitness.

For the final evolutionary layer, we incorporated pre-existing bioinformatics information, aligning 768 structural homologs of DraTnpB retrieved by Foldseek^15^. We computed per-residue conservation across the resulting multiple sequence alignment, designating the 87 positions conserved in at least half of the alignment as evolutionarily constrained (**Fig. 1e**). This set encapsulates residues that represent a putative evolutionarily conserved backbone of TnpB-like proteins.

The three layers proved substantially non-redundant, with only 7 residues nominated by all three layers (**Supplementary Fig. 2a**,**b**). We used the union of the three layers as the final sequence prompt, in which 151 residues were kept invariant while 257 of 408 positions (63%) were masked to be generated by ESM3. We additionally implemented Feynman–Kac inference-time steering^6^ (FK steering) to bias generation toward active sequences (**Methods**), using two independent reward functions that leveraged our internal DMS dataset **(Supplementary Fig. 3**) or an external DMS dataset of DraTnpB performed in yeast (**Supplementary Fig. 4 and 5**).

Across all conditions, we generated approximately 36,000 unique sequences. The resulting designs were divergent with a median of 57% identity to wild-type DraTnpB (approximately 176 substitutions) and 61% identity to one another, populating a broad but bounded region of sequence space that is distinct from the natural TnpB and Cas12f families (**Supplementary Fig. 6a)**. We filtered these candidates using a variety of metrics^16^ (**Methods**), selecting 44 for downstream experimental validation (**Supplementary Fig. 6b**,**c**).

### Experimental Characterization and Benchmarking of ACE Designs

We synthesized and tested nominated ACE designs for editing activity in human cells across a panel of loci that included the GFP-ON reporter locus used for the DMS screen and endogenous loci (**Fig. 2a**). Transfection of these designs into the GFP-ON reporter showed several (6/44) ACE constructs with significant editing activity, with some demonstrating equivalent levels to native DraTnpB (∼60% indel formation) (**Fig. 2b, Supplementary Fig. 7**).

**Figure 2.**
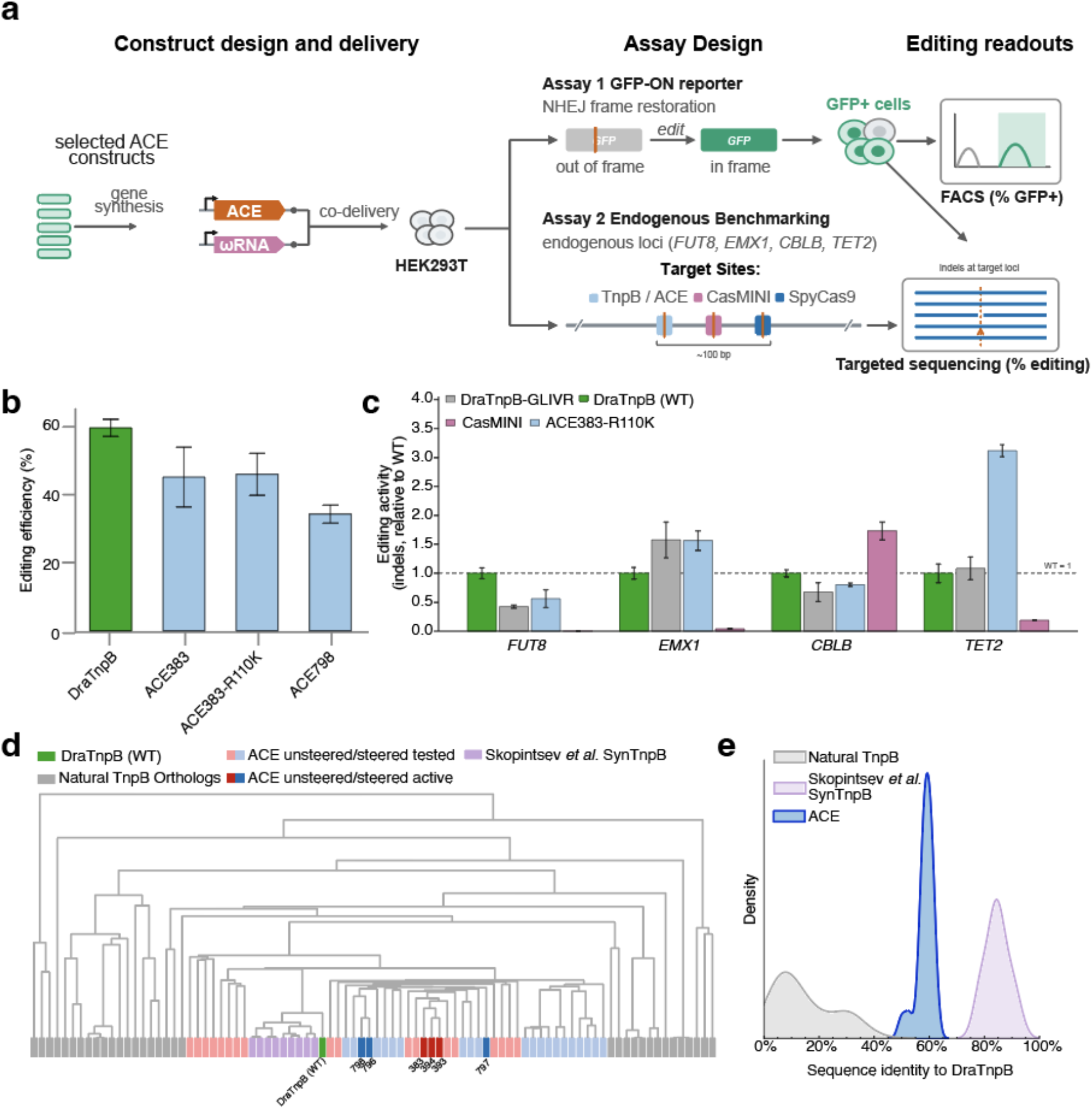
Characterization of the editing activity and sequence diversity of ACE constructs. **(a)** Construct and assay design. Selected ACE designs were synthesized and co-delivered with the cognate gRNA into HEK293T cells. Editing was assessed by the GFP-ON reporter or at selected endogenous loci. **(b)** Editing efficiency at the GFPon reporter for wild-type DraTnpB and select ACE constructs. **(c)** Editing activity at 4 selected endogenous loci, normalized to wild-type DraTnpB. **(d)** Dendrogram-based sequence similarity of AI-generated editors (using DraTnpB scaffold) and natural DraTnpB orthologs. Branch height is proportional to % sequence identity between two constructs in a multiple sequence alignment, with a lower branch height implying a higher degree of similarity. **(e)** Distribution of sequence identity (%) to DraTnpB for tested ACE and SynTnpB classes.

We found that the catalytically inactivating D191A mutation disabled editing for both native DraTnpB and ACE constructs. Additionally, our DMS screen revealed an insensitivity to mutations in the disordered C-terminal tail of DraTnpB (**Fig. 1b**). We verified that truncating DraTnpB to R380 maintained functional editing (**Supplementary Fig. 8**) and that the similarly truncated ACE was also active (**Supplementary Fig. 7b**). These results jointly show that functional insights can be carried over to the ACE context, implying a conserved mechanism of action. We also tested mutational variants that were previously described to enhance DraTnpB activity^11^. To our surprise, in our assay most of these mutations had no effect or were deleterious for both DraTnpB and ACE, except for an R110K mutation tested with the ACE383 construct (ACE383-R110K), which maintained equivalent or slightly superior editing relative to the base ACE383 editor.

We subsequently tested a panel of top ACE and DraTnpB variants along with SpyCas9 and CasMINI^17^ across four endogenous loci that contained overlapping target sites, allowing direct comparison of editing in the same genomic context^12,18^. We co-transfected plasmids encoding an editor and its cognate gRNA targeting each locus. SpyCas9 demonstrated consistently strong editing across loci (∼20%), while editing was more locus-dependent for the compact editors (**Fig. 2c and Supplementary Fig. 9**). CasMINI demonstrated low editing at all loci, except for *CBLB*, where it exceeded that of the TnpB systems. The TnpB editors generally maintained robust editing across the four loci tested, with ACE383-R110K showing similar levels of editing to wild-type DraTnpB, with activity relative to DraTnpB ranging from 0.57-fold at *FUT8* to 3.1-fold at *TET2*. Notably, ACE383-R110K matched or exceeded a previously described high-activity DraTnpB mutational variant (DraTnpB-GLIVR)^11^ at all four endogenous loci. The distribution of indel mutational outcomes from NHEJ at these endogenous loci for ACE383-R110K were indistinguishable from the native DraTnpB, again suggesting a similar nuclease mechanism for both ACE and native DraTnpB (**Supplementary Fig. 10**).

We subsequently analyzed the sequence properties of our tested designs relative to each other and pre-existing TnpBs. ACE and DraTnpB formed a cluster distinct from that of other natural TnpBs, confirming that DraTnpB is the closest relative to ACEs, as expected (**Fig. 2d**). In addition, active ACE designs predominantly cluster together, possibly indicating a shared sequence space for the active constructs. Intriguingly, the active ACE constructs were not trivially those closest to native DraTnpB in sequence space, demonstrating that they form a distinct cluster of active TnpB sequences. We also compared ACE sequences to those of a recent report generating novel DraTnpB-templated editors (SynTnpBs)^4^ using a related PLM, ESM-IF^19^. Consistent with their report, we found that SynTnpBs clustered much more strongly with DraTnpB and with each other, while ACEs showed significantly greater sequence diversity from DraTnpB (∼2-fold greater number of mutations) and from each other, even within functional subclusters (**Fig. 2d**,**e**). We further explored the nature of the amino acid changes in ACE relative to wild-type DraTnpB. In addition to being more numerous, we also found that substitutions within ACEs tended to be less chemically conservative than those within SynTnpB (**Supplementary Fig. 11**). Together, these results support that our approach produces highly diverse, novel, and programmable nucleases that maintain a high rate of activity.

## DISCUSSION

The editors produced here form a distinct clade of sequence space relative to native TnpBs, which suggests that this additional diversity can provide an alternative method to probe sequence determinants of nuclease function. Comparing active and inactive designs, we find residues that are strongly enriched in their respective classes (**Supplementary Fig. 12a)**. In particular, the most significantly enriched residue in our active class, which was found in all active ACE designs, was T228K. Notably, this residue was found to be an important residue in the SynTnpB structure due to its ability to mediate novel contacts with the gRNA scaffold^4^.

More generally, we noted a gain in positively charged residues within active ACEs that are predicted to interact with the gRNA (**Supplementary Fig. 12a**,**b**). This suggests a possible mechanism in which gRNA binding may be partially abrogated for synthetic TnpB proteins, thereby requiring compensatory mutations to enhance affinity to truncated versions of the DraTnpB gRNA commonly used within human cells. This is consistent with findings from a DMS screen of the DraTnpB gRNA, where the benefit of top gRNA hits did not transfer to mutational variants of DraTnpB^11^. Together these results suggest a sensitive interplay between the gRNA and TnpB mutational variants, which may explain the discrepancy between the reported gain in function for DraTnpB mutational variants and the results in this work, which derive from an ACE system that contained a gRNA scaffold that was further truncated.

We hypothesized that the addition of a functional layer to structural and evolutionary layers within a prompt would be critical for recapitulating editing function. We performed an *in silico* analysis to explore how each information layer contributes to the sequences generated (**Supplementary Fig. 13**). By removing the information from individual layers and prompting ESM3, we find that the set using all three layers resulted in sequences that cluster distinctly from those prompted with the reduced information sets and more closely to native TnpBs. This analysis supports that that the information layers are complementary and combine to form a unique phenotype that consists of active genome editors within a subset of those tested. Notably, randomly fixing the same number of residues is also insufficient to reconstitute ACE-like constructs. Whether this ACE-cluster is enriched for active constructs stands as an interesting area of continued exploration.

A major distinguishing feature of our method is the significant increase in diversity in our generated and active constructs (∼40% divergence from the native DraTnpB) relative to prior reports^2–4^, which classifies them closer to the realm of related orthologs rather than sequence variants (**Supplementary Fig. 14**). Additionally, our method is computationally efficient, with a hit rate per generated construct 10-1000x higher compared to other approaches, while also comparing favorably on hit rate per construct tested and scale of downstream experimental screening required to identify an active construct. This suggests that the method may be useful to effectively navigate the vast sequence space for other protein engineering applications.

In addition to the methodological improvements of the generative approach, the generated editors described here demonstrate a favorable profile for use in therapeutic applications, with greater utility than SpyCas9 for cargo-constrained delivery applications and an overall activity level that is equivalent or superior compared to existing compact editors. In addition, the ACE383-R110K construct had the lowest predicted immunogenicity (**Methods**) relative to the benchmark editors, which is consistent with a previous report^2^ suggesting that AI-generated editors tend to have a lower degree of immunogenicity relative to editors that are naturally occurring (**Supplementary Fig. 15**). To our knowledge, the generated set of ACEs also contains, at 380 amino acids, the smallest synthetic RNA-guided nuclease described to date.

This work describes a framework for creating highly diverse, novel proteins that can recapitulate complex scaffolding and enzymatic functions. Analogously to prompt engineering of large LMs (LLMs), generating more sophisticated prompts and steering algorithms for PLMs will significantly improve their performance for bespoke use cases without requiring laborious fine-tuning. We show that by combining efficient data collection and bioinformatics approaches, it is possible to create highly effective prompts to harness an off-the-shelf foundational PLM to generate functional proteins in sequence space that are distinct from naturally evolved proteins. The resulting proteins serve as useful substrates for determining protein sequence-function relationships, which will serve as a critical step for designing both optimized and novel functions for downstream biotechnological and therapeutic applications.

## ACKNOWLEDGEMENTS

The authors thank Dr. Monte Winslow, Dr. Doug Fowler, and Dr. Le Cong for comments on this study and manuscript. The authors also thank the NVIDIA Inception Program for a computational grant. Research reported in this publication was supported, in part, by the National Institute on Aging of the National Institute of Health under award number R43AF088777 (to N.W.H).

## MATERIALS AND METHODS

### Cell line and cell culture

HEK293T cell lines were cultured in DMEM, high glucose supplemented with 10% fetal bovine serum, 100 U/mL penicillin and 100 mg/mL streptomycin at 37 °C with 5% CO_2_.

### Lentiviral preparation and delivery

Lentivirus was prepared by co-transfecting transfer plasmids with VSV-G envelope and Delta-Vpr packaging plasmids into HEK-293T cells using Lipofectamine 3000 reagent (ThermoFisher). Supernatant was harvested 48 h and 72 h after transfection. HEK293T cells were transduced at high-MOI with the gRNA and GFP-ON reporter components followed by selection to recover a pure population that expressed each component.

For the preparation of large-scale lentivirus for the pooled deep mutational scanning assay, several modifications were made to the small-scale protocol. First, lentivirus was concentrated by 10X using PEG-it virus precipitation solution (System Biosciences) according to the manufacturer’s instructions. Concentrated virus was then used for a functional titer in which a concentration consistent with an MOI of ∼0.3 was identified. Subsequently, cells stably expressing the reporter components were transduced at a scale to achieve ∼1000X cells/ISDra2 variant after puromycin selection of the pooled library component.

### GFP-ON Reporter Construction

A reporter construct containing the GFP-ON system was constructed with a synthetic target site flanked upstream by a blasticidin resistance gene and downstream by an out-of-frame EGFP gene (via NEB HiFi Assembly). The frame disorientation was designed such that all 3n+2 length indel mutations would reconstitute the blasticidin-2A-GFP reading frame, leading to the activation of EGFP expression. The construct bearing the gRNA scaffold sequence and guide targeting the GFP-ON reporter was assembled through PacI/NdeI double digestion of a lentiviral construct and ligation of a gene fragment containing the omegaRNA and guide sequences downstream of a human U6 promoter. These constructs were packaged and lentivirally introduced.

### DraTnpB DMS design and implementation

All possible single amino acid substitutions were generated by staging NNK codons across the length of the DraTnpB protein body. A custom python script was used to create 5 individual tiled regions that were within the length requirements for high-throughput oligonucleotide synthesis (<300 nucleotides). These tiles were cloned with 4 native-sequence tiles into a lentiviral plasmid, inserting at an EF-1a core promoter and upstream of a 2A peptide /puromycin resistance cassette, forming 5 sub-libraries, each containing a region of single-amino acid substitutions that were then combined to form the final library.

### Next-generation sequencing analysis workflow and quantification of variant fitness effect

Primers were designed to amplify each tiled region separately and contained Illumina adapters to enable flow-cell clustering and next-generation sequencing. Reads were processed using a custom pipeline. Briefly, reads were trimmed and merged using publicly available software (cutadapt4.4 and FLASh v1.2.11). Next, the tiled region was aligned using the Needleman-Wunsch algorithm to a reference sequence to discard reads that contain problematic indel mutations or chimeras. The resulting length-matched nucleotide sequence was then translated into an amino acid sequence and parsed to call variants.

A data frame was constructed that contained the abundance of variants within each sample to enable input into the dmsTools2^20^ differential selection workflow using default parameters across a range of read thresholds. Prior to the dmsTools2 workflow, the data frame was manually curated to remove spurious, lowly represented, variant calls (i.e. multiple amino acid variants) that were not expected to occur through the NNK-based mutagenesis strategy.

### Prompt engineering and design generation without steering

A local (3DiAA) FoldSeek search was conducted and yielded 9101 hits across multiple databases (BFVD-1179, afdb-proteome-218, afdb-swissprot-363, afdb50-3000, bfmd-218, cath50-465, gmgcl_id-218, mgnify_esm30-3000, pdb100-440). The hits from the search were filtered to unique sequences and to retain those with an alignment probability of 1, an e-value <= 1e-20, and alignment length that was +/-10% of the query sequence length, which yielded 768 protein sequences. Clustal-Omega was used to perform a global alignment from which residues were selected due to their degree of conservation across the set of protein sequences (>50%). The final sequence prompt was created by taking a union of all the approaches which yielded a prompt with 257/408 (62%) of the residues masked. We computationally generated a total of 31,500 designs using an A100 (NVIDIA) GPU, using sequence-only prompting and prompting using combined sequence and structure tracks. We retained 95 designs whose pTM >= 0.90 and global_RMSD was <= 3, selecting 21 for downstream assessment.

### Reward function using external DMS data

A multilayer perceptron (MLP) was trained to predict variance-normalized deep mutational scanning (DMS) enrichment scores from an external DMS dataset (single and combinatorial variant libraries)^11^. Each candidate sequence was embedded using ESM2 650M, and the mean-pooled 1280-dimensional final hidden-state representation was standardized and passed through a feedforward network (1280→512→256→128→1, with LayerNorm, GELU activation, and dropout = 0.2 at each hidden layer) to yield a scalar predicted fitness score. The model was trained by minimizing mean-squared error against measured enrichment scores, using the AdamW optimizer (learning rate = 1e-3, weight decay = 1e-4) with cosine-annealed learning-rate decay and early stopping (patience = 10 epochs) on a held-out validation split (random 10% subset of full data). Because the reward model was queried on partially masked sequences during FK decoding, training data were augmented with copies of each sequence masked at a randomly sampled fraction of variable positions (uniform on [0, 1]) while retaining the original fitness label.

### Reward function using internal DMS data

A second reward function used DMS data from the screen presented within this study. Briefly, every possible single substitution in the 408-residue DraTnpB parent was scored by the mean of a per-substitution Bayesian posterior, yielding a 408 × 20 reward table. The prior was an ESM2 masked-marginal zero-shot score, z = log P(mut) − log P(wt), linearly calibrated into DMS units by ordinary least squares against the measured variants: μ_0_ = a·z + b, with a = 0.1539 and b = 0.1116 (N = 3,993; R^2^ = 0.173). The prior standard deviation, σ_0_ = 1.107, was taken as the root-mean-square residual of that fit and assigned uniformly to every cell. The likelihood was the replicate-aggregated DMS measurement *d* (mutdiffsel; variants retained at ≥ 2 batches) together with its batch-to-batch standard deviation *σ*_*d*_, clamped to [0.05, 5.0]. Writing precision as τ = 1/σ^2^, each measured cell was updated by the conjugate normal–normal rule μ = (τ_0_μ_0_ + τ_d_·*d*)/(τ_0_ + τ_d_) and σ = (τ_0_ + τ_d_)^(−1/2); cells without a measurement retained the prior unchanged (3,759 of 7,752 non-wild-type cells), and wild-type identities were fixed at μ = σ = 0.

### Feynman-Kac (FK) steered generation

Candidate sequences were generated using an FK particle-steering procedure applied to the iterative masked-token decoding trajectory of ESM3 (esm3_sm_open_v1). Briefly, a population of K = 50 particles was initialized as identical copies of the parent sequence with the same positions masked/unmasked as specified in the main text. An additional 21 positions were fixed to wildtype at the start of generation. These positions matched the 21 positions that, during generation without steering, were originally masked but regenerated back to the wildtype residue identity. At each decoding step, ESM3 was used to compute the conditional token distribution for all particles, and for each particle residues were unmasked by sampling directly from the ESM3 predictive distribution: *P*(*x*_*i ∈ masked*_|*x*), where *x*_*i ∈ masked*_ denotes the masked residues in *x*, and *x* denotes the full sequence at the current denoising step (including masked positions). Masked positions are decoded according to ESM confidence per position, max 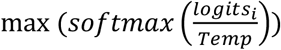 or the model’s certainty about the top residue guess at each masked position. A fixed number of positions were unmasked at each decoding step. The sampling temperature was fixed to 0.7 across all runs. After each unmasking step, every particle’s partial sequence was scored using a fitness-prediction reward model (see below), and importance weights were computed from the change in predicted score between consecutive steps (scaled by an additional temperature parameter τ) and normalized by softmax. Particles were resampled using a systematic resampling scheme whenever the effective sample size 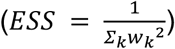 fell below 50% of the population, replacing low-weight particles with duplicates of high-weight particles.

In addition, the reward table generated using the Bayesian reward framework derived from the internal DMS data developed in this study was was applied by Feynman–Kac particle steering: K = 50 particles were decoded in parallel over 10 iterative unmasking steps, sampling at every step from the unmodified ESM3 distribution, and each particle was scored by summing μ − βσ over its decoded positions (uncertainty penalty β = 0.5). Particles were reweighted by the score increment between successive steps, log w_t_ ∝ (r_t_ − r_t−1_)/τ, and resampled whenever the effective sample size fell below K/2, so the reward acts through population-level selection; τ was swept over {0.1, 0.5, 1.0, 5.0, 10.0} with 10 independent campaigns per setting (500 candidates each).

### Design testing within human cells

The nominated designs were synthesized as eBlocks (IDT) and assembled into a plasmid backbone using HiFi Assembly (NEB) downstream of EF1 αand upstream of a 2A-Puromycin resistance cassette. HEK293T cells bearing the GFP-ON reporter system and an gRNA targeting it were transfected with plasmids bearing ACE designs using Lipofectamine 3000. Briefly, 5e4 cells were plated 24 hours prior to transfection and transfected with 500 ng of plasmid with puromycin selection (2 µg/mL) optionally provided at days 1-3 post-transfection. Cell samples were analyzed for editing activity at 3-10 days post-transfection. Editing was quantified using FACS-based measurement of GFP levels, as well as with targeted amplification of the target locus followed by quantification of indel accumulation using CRISPResso2 on data collected through Oxford Nanopore Technologies (ONT) sequencing.

### Benchmarking nuclease activity across endogenous and synthetic loci

All transfections were performed with Lipofectamine 3000 (Invitrogen) at a ratio of 1 µl of reagent per microgram of plasmid and per 5-µl volume of Opti-MEM Reduced Serum Media (Gibco). On the day of transfections cells were seeded into 96well plates at a concentration of 4e5 cells/mL. For the gene editing experiments targeting endogenous genomic loci (Fig. 2c), 250 ng of nuclease plasmids and 250 ng of gRNA plasmids were transfected into HEK293T cells. The transfected cells were selected through puromycin 24hrs after transfection and genomic DNA was extracted using QuickExtract (LGC, Biosearch Technologies) 6 days after transfection for deep sequencing analyses.

### Immunogenicity prediction

MHC binding was predicted for effector proteins using NetMHCpan 4.2c for class I and NetMHCIIpan 4.3j for class II, scanning each sequence for all 8 to 11 residue peptides against a 27 allele class I panel and all 15 residue peptides against a 26 allele class II panel, with binding affinity prediction enabled. Strong binders were classified as %Rank ≤ 0.5 and weak binders as %Rank ≤ 2.0 using standard cutoffs, and binders were counted per protein and allele pair. The strong binder scores for both the classes were summed to give a single predicted immunogenicity value per effector.

