## Supplementary Information for "Efficient exploration of sequence space enables rapid generation of functional genome editors"

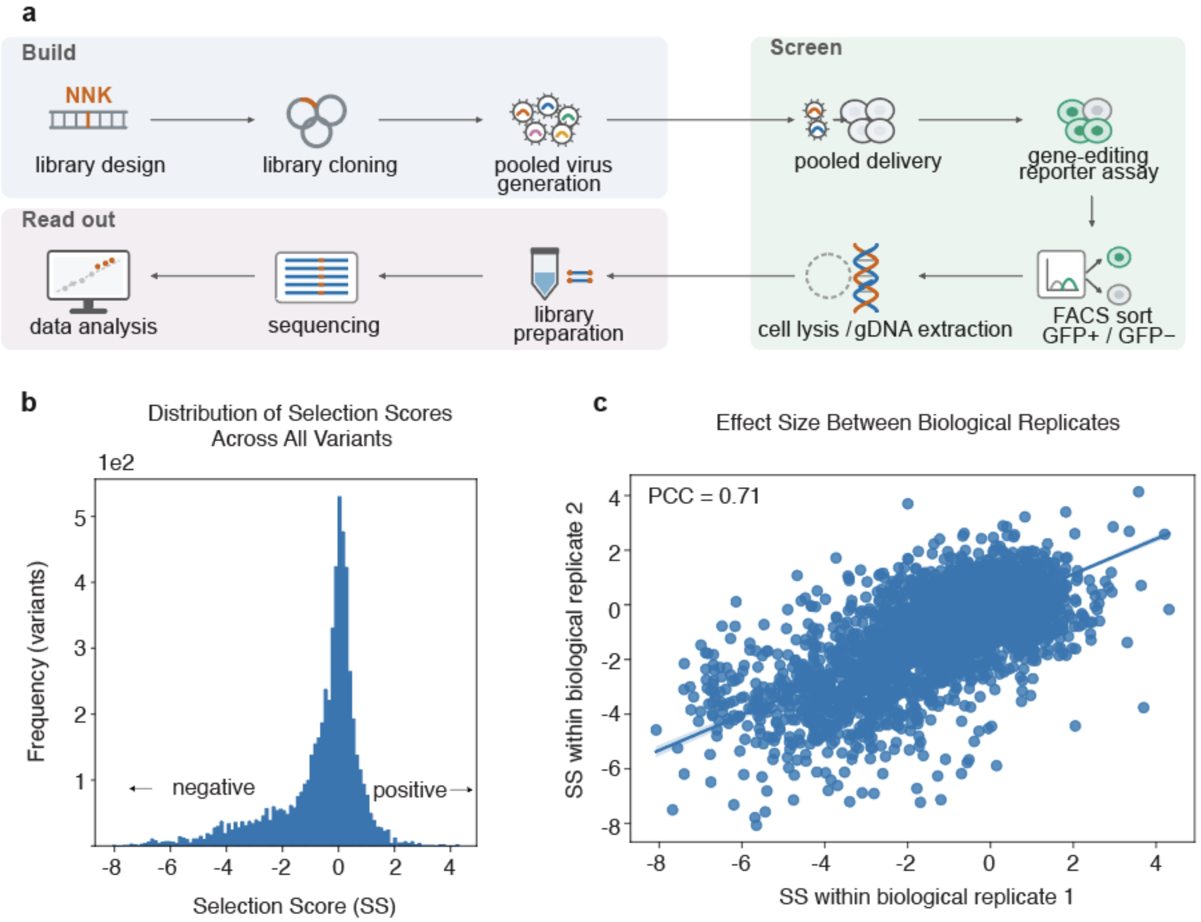


**Supplementary Figure 1. Development of a pooled screening system for deep mutational scanning of the DraTnpB nuclease. (a)** Schematic of the pooled screen experimental design in which edited HEK293T cells express GFP and are enriched using FACS followed by NGS-based quantification of variant identity. **(b)** Distribution of selection scores (SS), defined as the mean of the mutdiffsel (**Methods**) across each experimental batch for all tested variants. **(c)** Correlation of SS by residue across biological replicates.

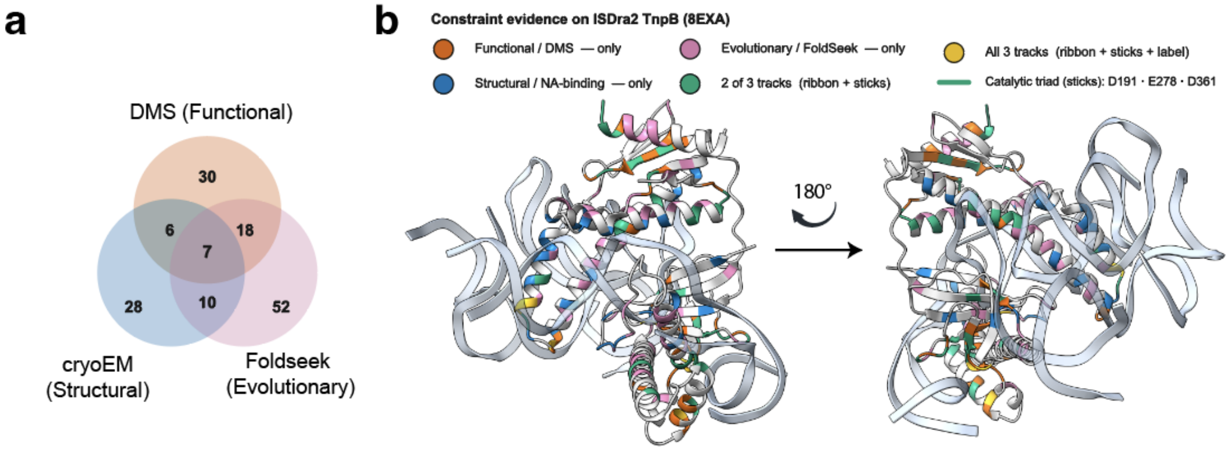


**Supplementary Figure 2. Prompt design to generate the ACE design space. (a)** Overlap among the residues nominated by the three evidence layers used to construct the prompt. **(b)** Mapping of residues nominated by each layer onto the DraTnpB structure (PDB 8EXA), colored by nomination source.

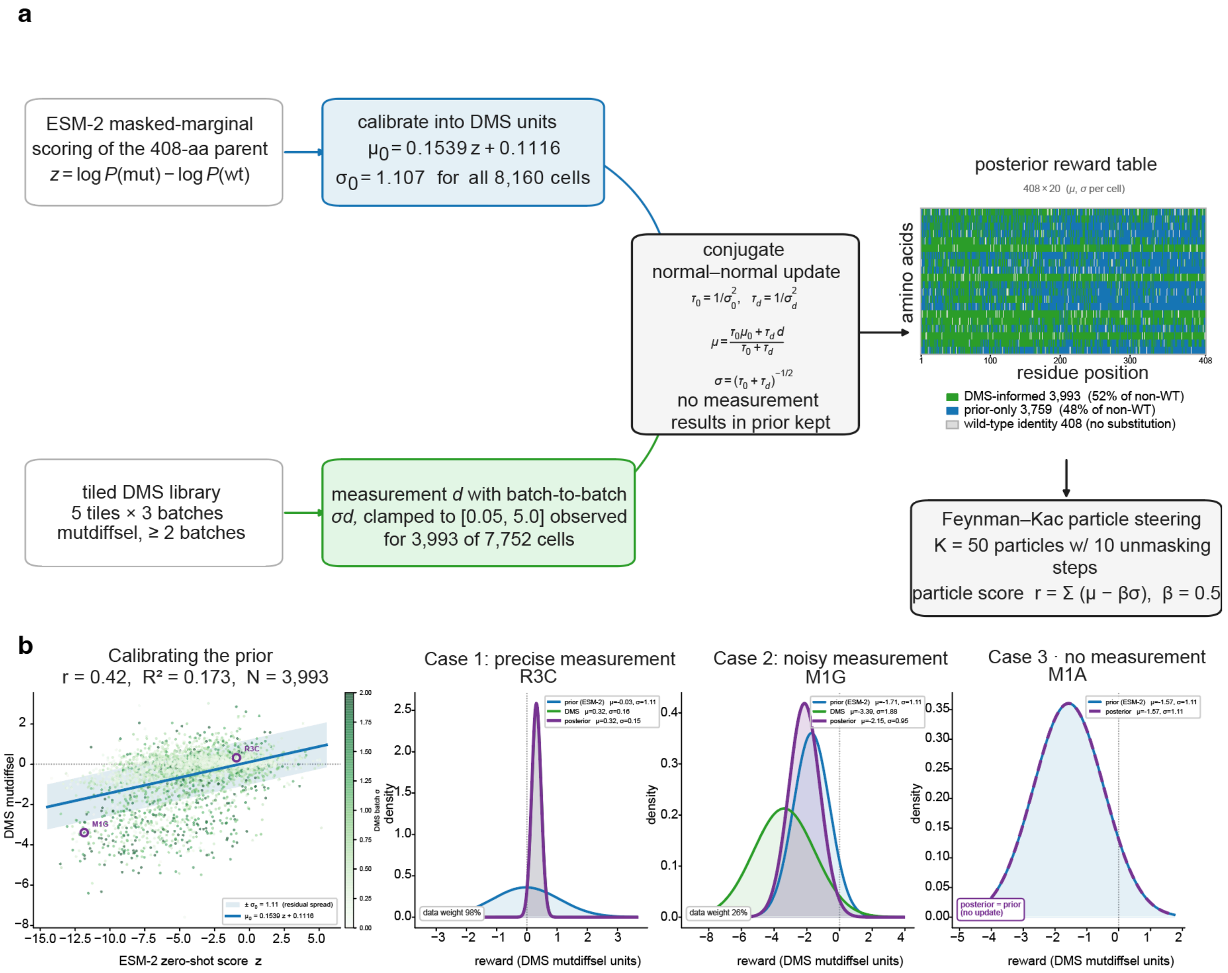


**Supplementary Figure 3. Feynman-Kac (FK) steering of ESM3 using a reward function based on an internal DMS dataset. (a)** A calibrated ESM-2 zero-shot prior and the tiled DMS measurements are combined by a conjugate normal–normal update into a per-cell posterior (mean μ, s.d. σ) covering all 408 × 20 substitutions, which then steers generation by Feynman–Kac particle selection in which particles are sampled from the unmodified ESM-3 distribution and reweighted by their accumulated reward, rather than by modifying logits (**Methods**). The grid gives each cell's provenance: DMS-informed (green, n = 3,993), prior-only (blue, n = 3,759; 48.5% of non-wild-type cells) and wild-type identity (grey, n = 408). **(b)** Left, prior calibration: DMS mutdiffsel against ESM-2 zero-shot score (N = 3,993; r = 0.42, $R^2$ = 0.173), with the fitted line and the $\pm\sigma_0$ = 1.11 residual band that sets the prior s.d.; circles mark the two exemplars shown at right. Right, the update for three representative cells (prior, blue; DMS, green; posterior, purple): a precise measurement dominates the prior (R3C, 98% data weight), a measurement noisier than the prior does not (M1G, 26%), and an unmeasured cell retains it unchanged (M1A, dashed). Curves are unit-area probability densities.

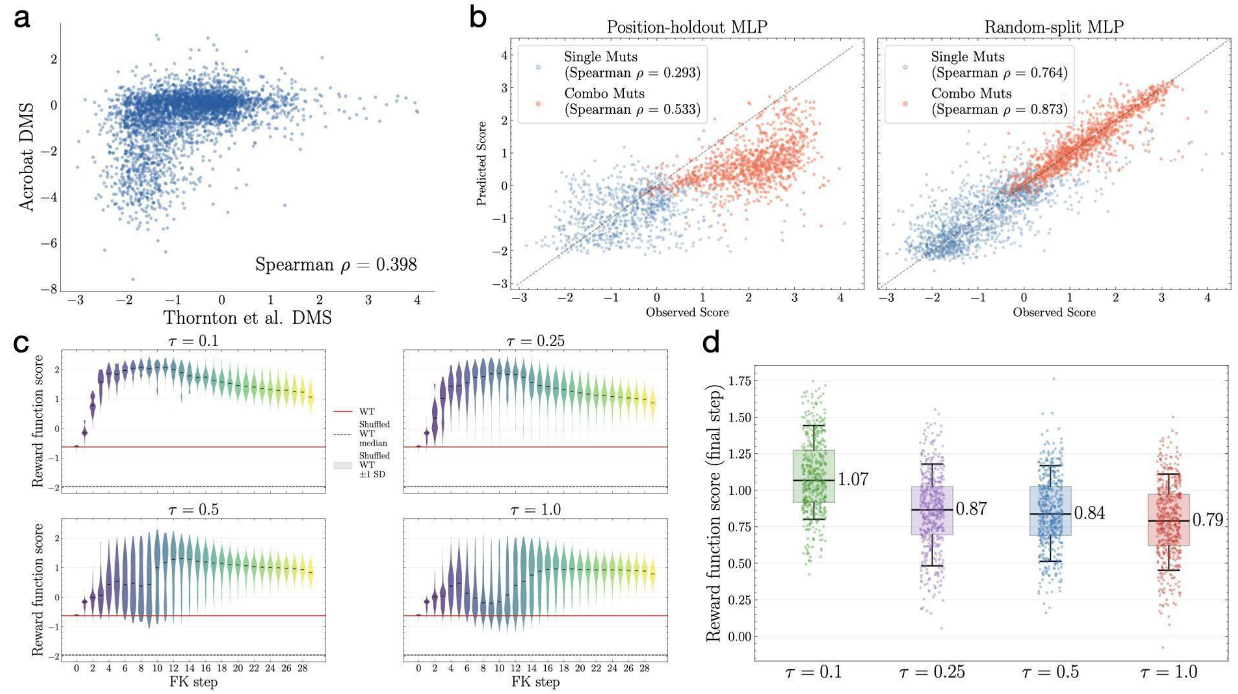


**Supplementary Figure 4. Feynman-Kac (FK) steering of ESM3 using a reward function on an external dataset. (a)** Correlation of effect sizes between Thornton et al. and Acrobat DMS of the DraTnpB protein. (**b)** Predictive power (Spearman correlation) of an MLP model trained on the Thornton et al. DMS across two different test-train split regimes: a position-based split, where all mutations within a position are either in training *or* testing, and a random split of the data. Correlation is reported separately for single versus combinatorial mutations. **(c)** Distribution of reward function scores across 30 ESM3 denoising steps and four distinct τ values (inversely proportional to resampling strength, see **Methods**). The score of the DraTnpB sequence under the reward function is shown in red. The DraTnpB was randomly shuffled and scored 10 times, and the median score and standard deviation is reported in grey. **(d)** The distribution of reward function across all steered candidates, reported within the τ value used for generation. The median score is reported.

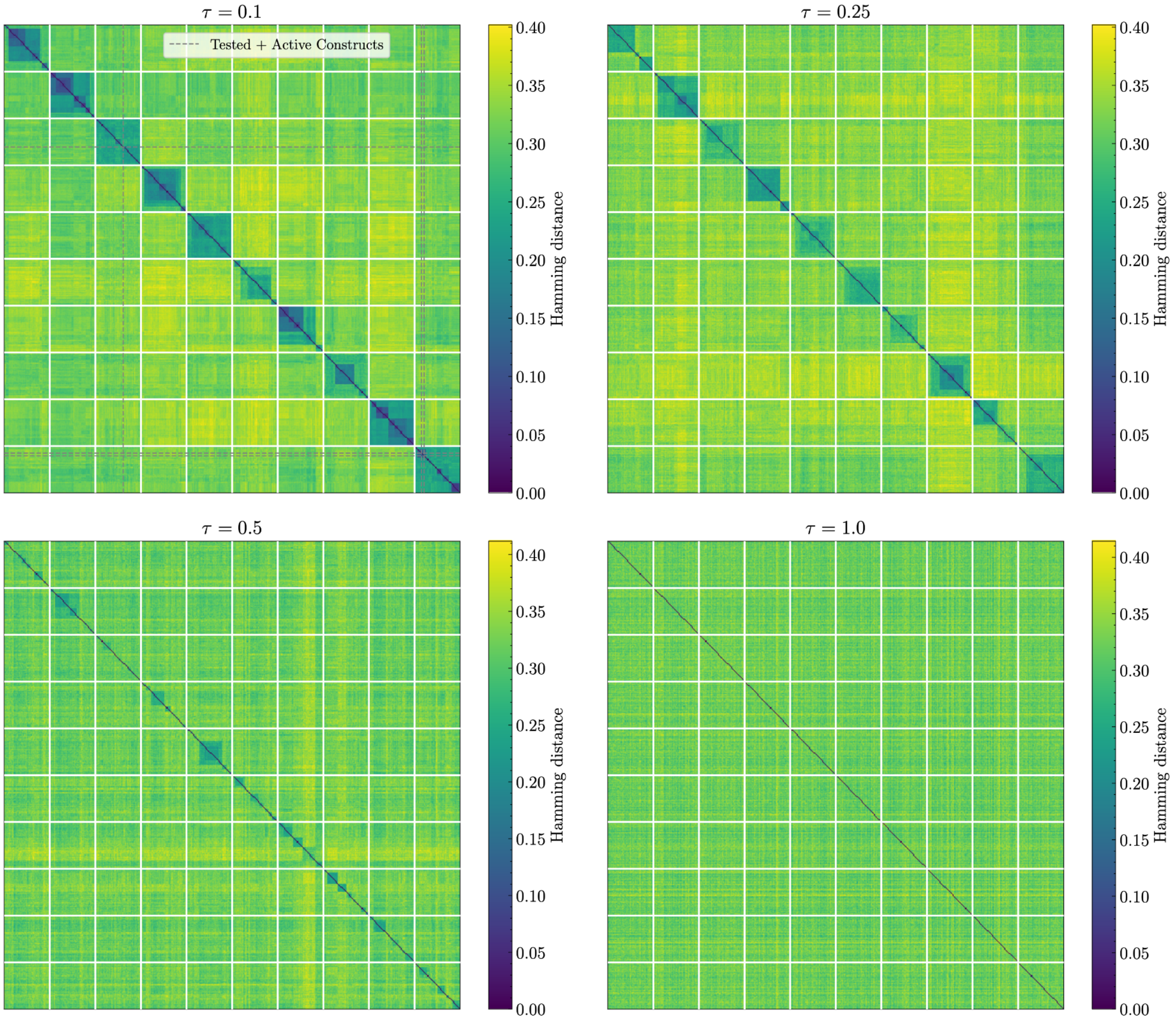


**Supplementary Figure 5. Sequence similarity of generated editors across different design campaigns and steering hyperparameters.** Heatmap depicting the pairwise Hamming distance between all constructs generated via FK steering with the external dataset reward function. FK steering was used to generate 2,000 unique constructs across four levels of τ. Within each τ, 10 design campaigns were run, with 50 particles per campaign. Constructs found to be active in downstream experiments are denoted by dashed lines.

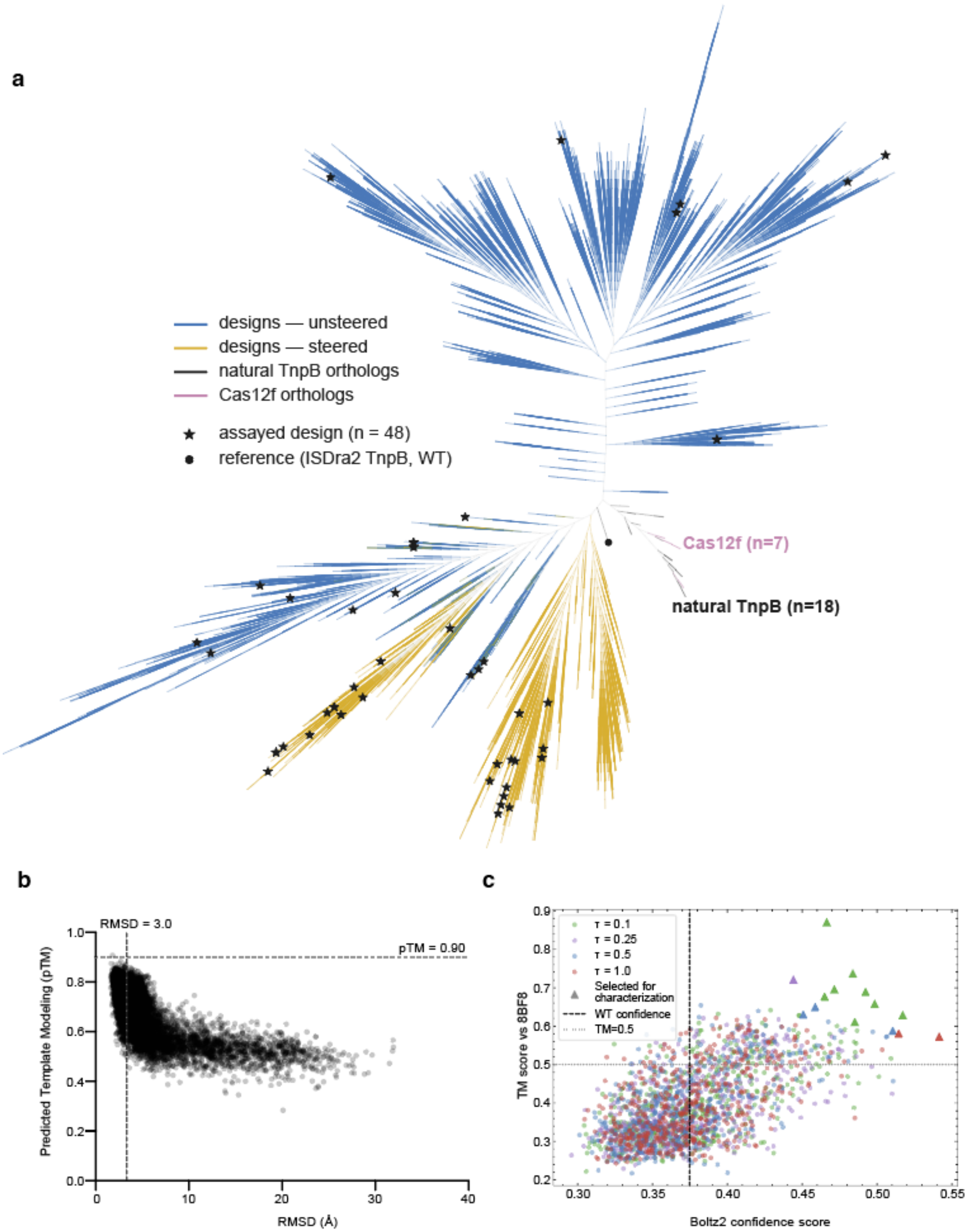


**Supplementary Figure 6. Diversity and filtering of generated constructs for experimental validation.**

**(a)** Cladogram of the design space, showing unsteered and steered designs alongside natural TnpB orthologs and Cas12f orthologs. Stars mark the 48 designs selected for experimental assessment. The filled circle marks the DraTnpB reference. **(b)** Unsteered constructs were scored against structural similarity to the DraTnpB structure, as quantified by RMSD, and pTM. The top 95 candidates whose pTM >= 0.90 or RMSD <= 3 were retained and rank-ordered for experimental testing **(c)** Steered constructs were co-folded (via Boltz2) with the gRNA from the 8BF8 structure (DraTnpB-gRNA complex) scored on the Boltz2 confidence of the predicted structure and the similarity (TM score) to the 8BF8 structure. The Boltz2 confidence score of DraTnpB with the 8BF8 reRNA is denoted by a vertical dashed line. The confidence score and TM score were summed, and the top constructs were selected for downstream experimental characterization. The plot shows a representative example corresponding to the constructs generated using the first (external) reward function.

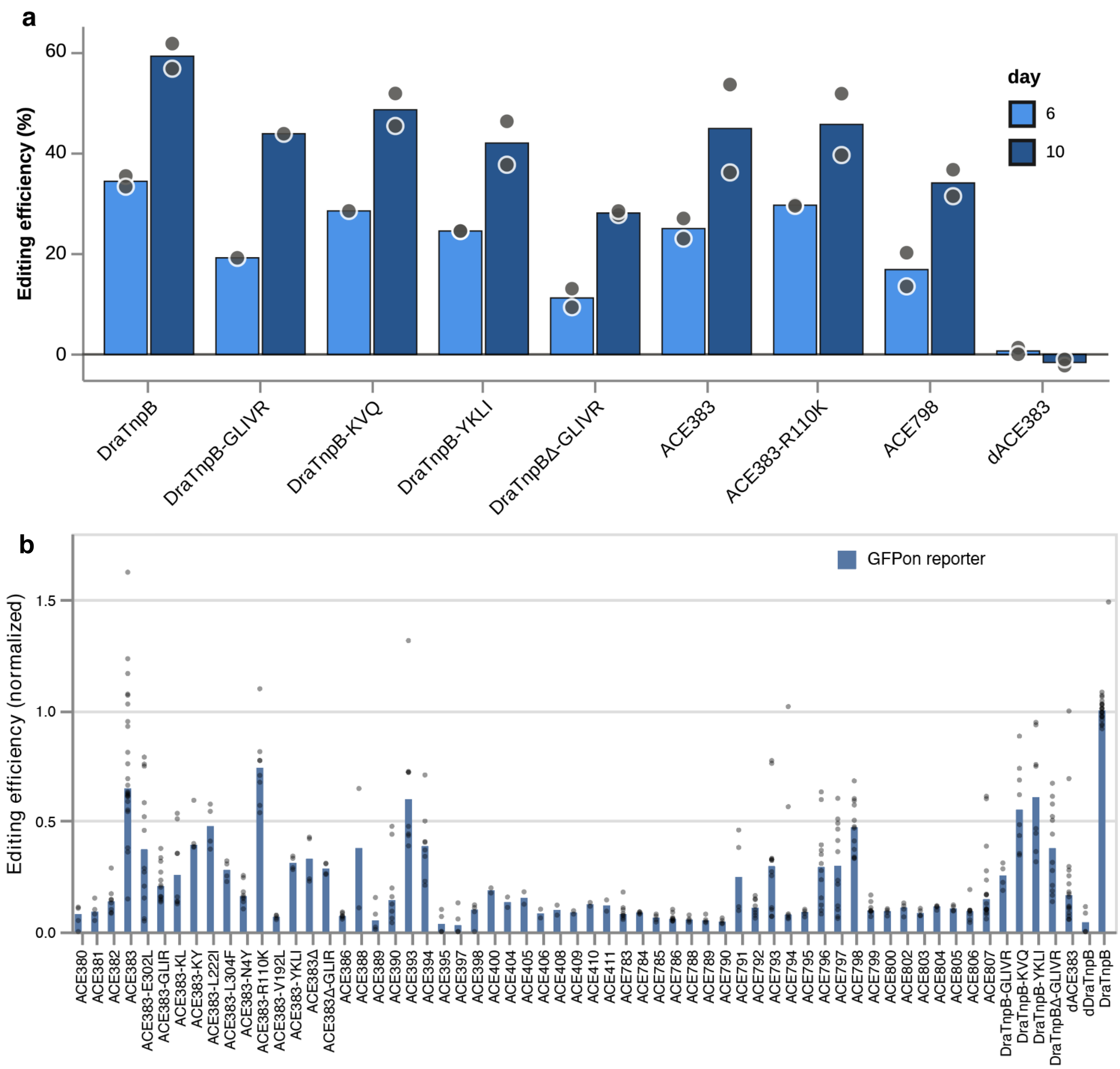


**Supplementary Figure 7. Editing activity of ACE designs at the GFP-ON reporter. (a)** Editing efficiency 6 and 10 days after transfection for wild-type DraTnpB, three previously reported engineered DraTnpB variants (GLIVR, KVQ, YKLI), a truncated derivative (DraTnpBΔ-GLIVR), ACE383, ACE383-R110K, ACE798, and the catalytically inactive control dACE383. Editing efficiency is measured by nanopore sequencing and is background-corrected by subtracting basal editing rate called by untransfected cells. Bars, mean; points, individual replicates; n = 2. **(b)** Editing efficiency across all assayed constructs measured in the GFP-ON reporter cells by flow cytometry, normalized to wild-type DraTnpB. Bars: means; points: individual replicates.

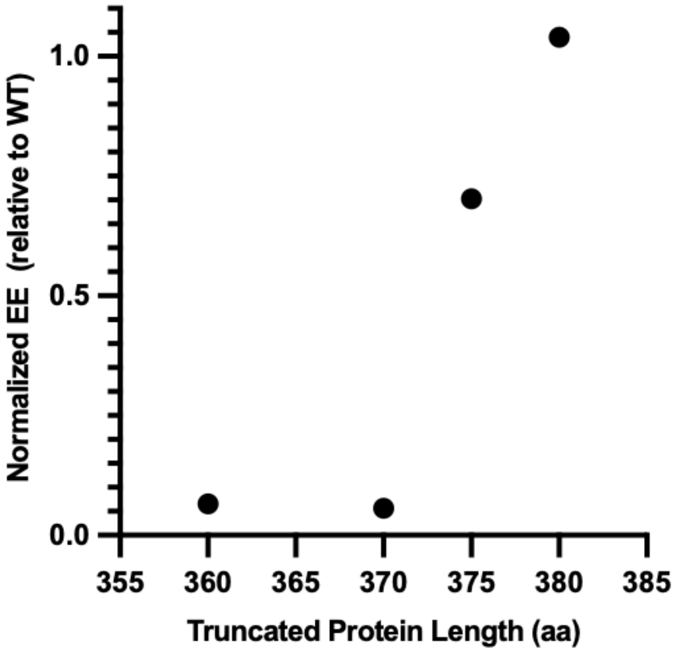


**Supplementary Figure 8. Performance of DraTnpB truncations.** Editing efficiency (normalized to full-length DraTnpB) of C-terminal truncations of wild-type DraTnpB, with the last residue of each construct indicated on the x-axis.

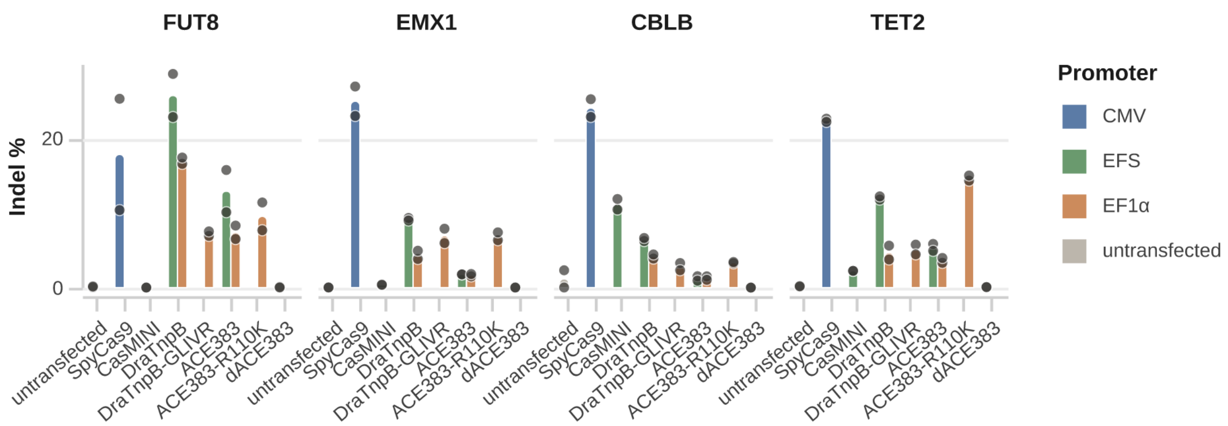


**Supplementary Figure 9. Editing at endogenous loci across effectors.** Indel frequency at *FUT8*, *EMX1*, *CBLB* and *TET2* for untransfected cells, SpyCas9, CasMINI, wild-type DraTnpB, DraTnpB-GLIVR, ACE383, ACE383-R110K and the catalytically inactive control dACE383. Bar colour denotes the promoter driving each effector (CMV, EFS or EF1α) Bars, mean; points, individual replicates; n = 2.

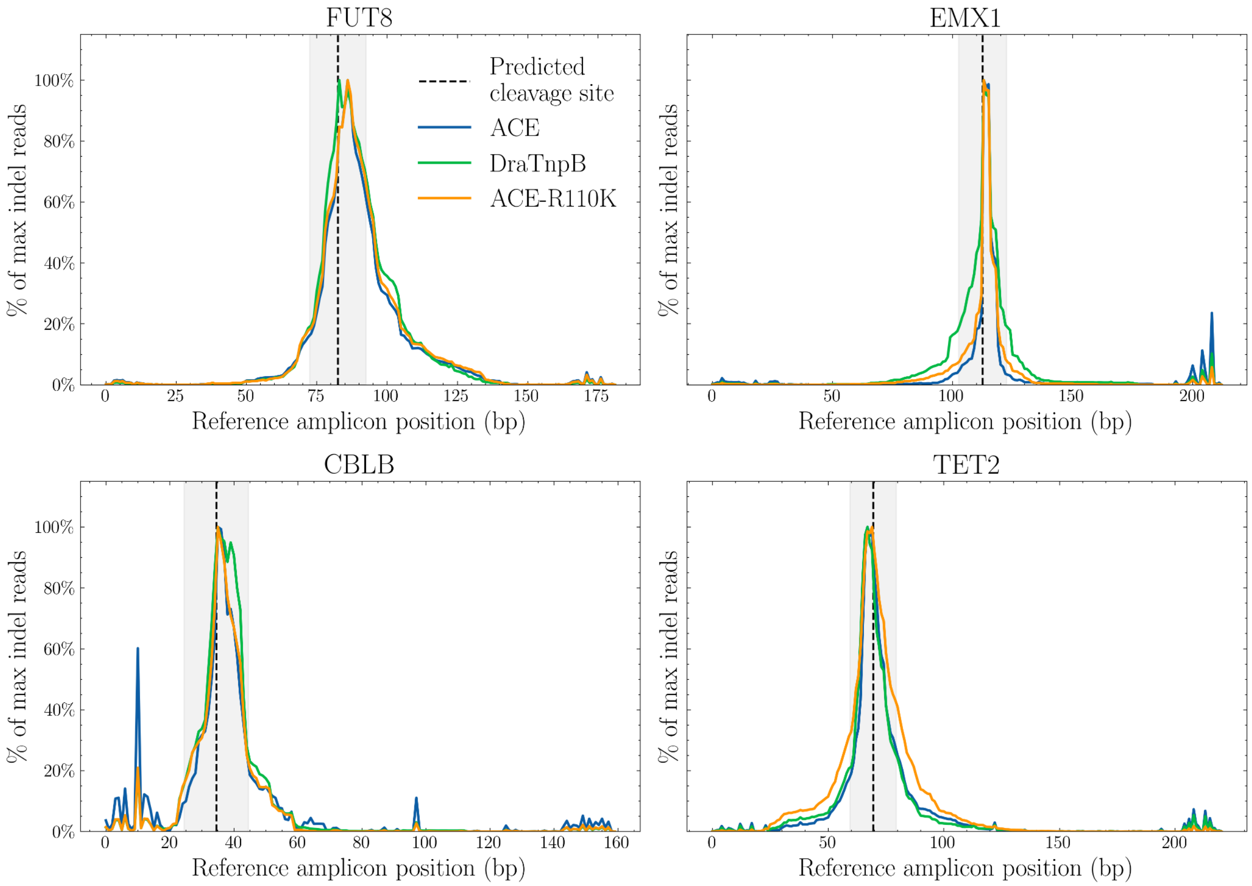


**Supplementary Figure 10. Indel profiles of DraTnpB and the generated ACE constructs at endogenous targets.** For each target, the percentage of indel-containing reads (insertions and deletions, pooled across replicates; substitutions excluded) is plotted against position along the amplicon. Each curve is normalized to that sample's own peak indel-read count, so all curves reach 100% at their maximum regardless of absolute editing efficiency. The predicted cut site and quantification window are highlighted.

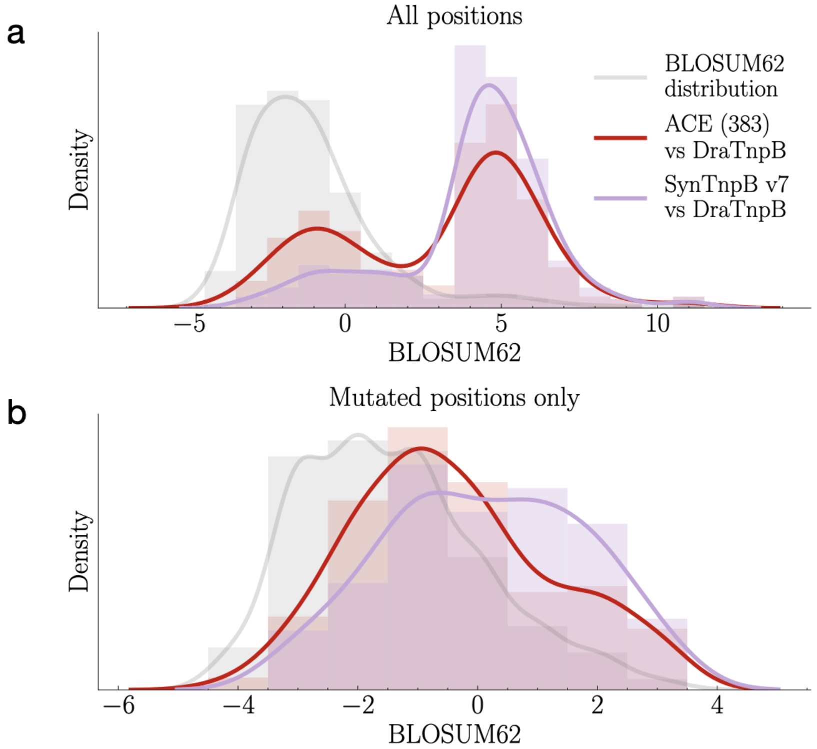


**Supplementary Figure 11. Comparison of substitution conservation of ACE383 and SynTnpB v7 against DraTnpB.** Distribution of conservation scores for substitutions of ACE383 (red) and SynTnpB v7 (purple) relative to wild-type DraTnpB, as scored by the BLOSUM62 matrix. The background distribution of BLOSUM62 scores (including the matrix diagonal) is included in gray for reference. Kernel densities are provided as a visual aid. **(a)** Comparison of all residues against DraTnpB. $p$=1.0e-8 for t-test of distributions between ACE383 and SynTnpB. **(b)** Comparison restricted to sites that are mutated relative to wildtype for ACE (n=165 positions) and SynTnpB v7 (n=85). $p$=0.014.

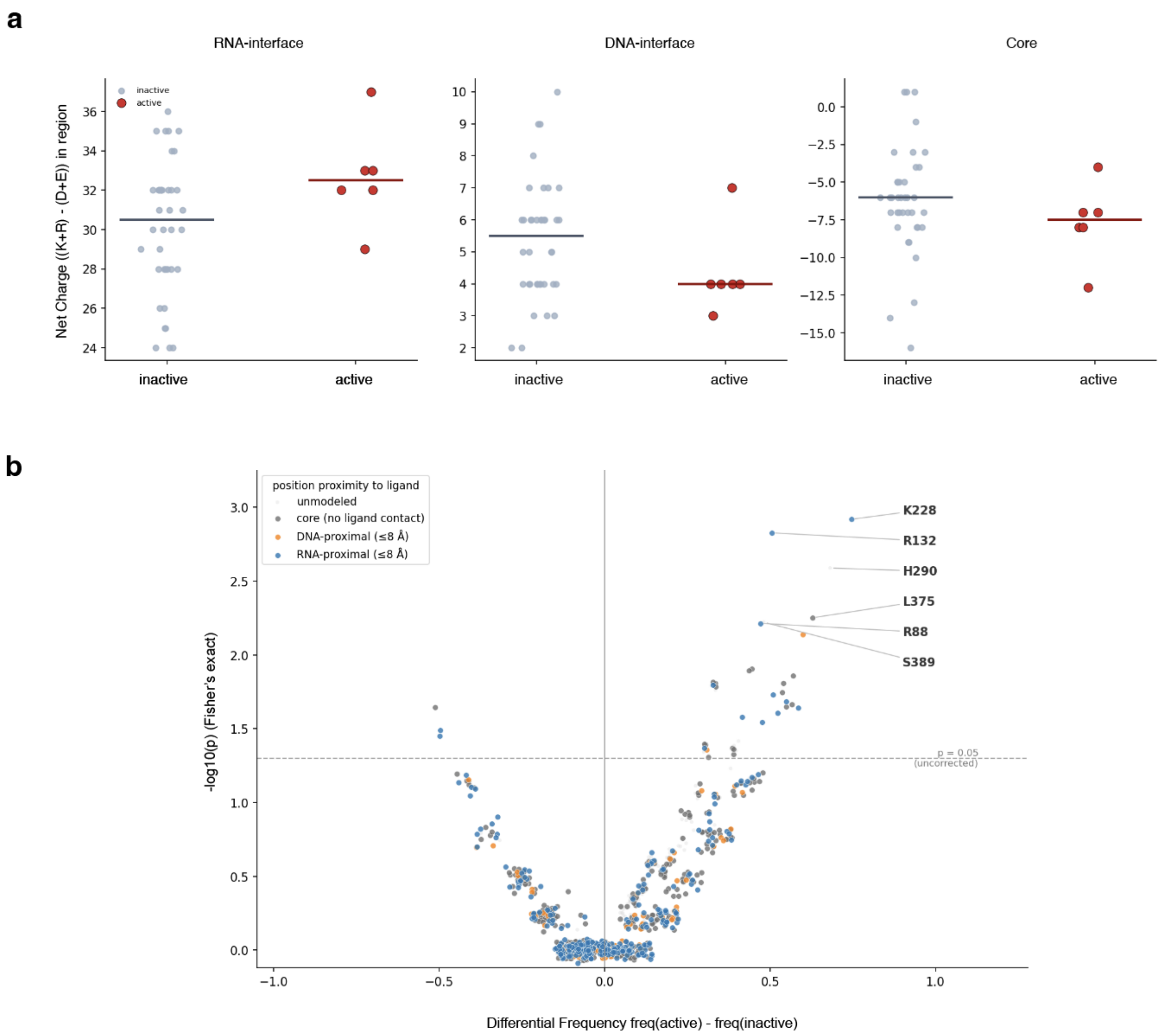


**Supplementary Figure 12. Sequence features distinguishing active from inactive ACE designs.** (**a**) Net charge ((K+R) − (D+E)) computed within the RNA-interface, DNA-interface and core regions for active (red) and inactive (grey) designs; horizontal bars, group medians. (**b**) Differential residue frequency between active and inactive designs. Each point is a specific residue; the x-axis gives the difference in residue frequency between the active and inactive sets and the y-axis the $-\log_{10} p$ (Fisher's exact test, uncorrected). Points are coloured by proximity to bound ligand in the reference structure.

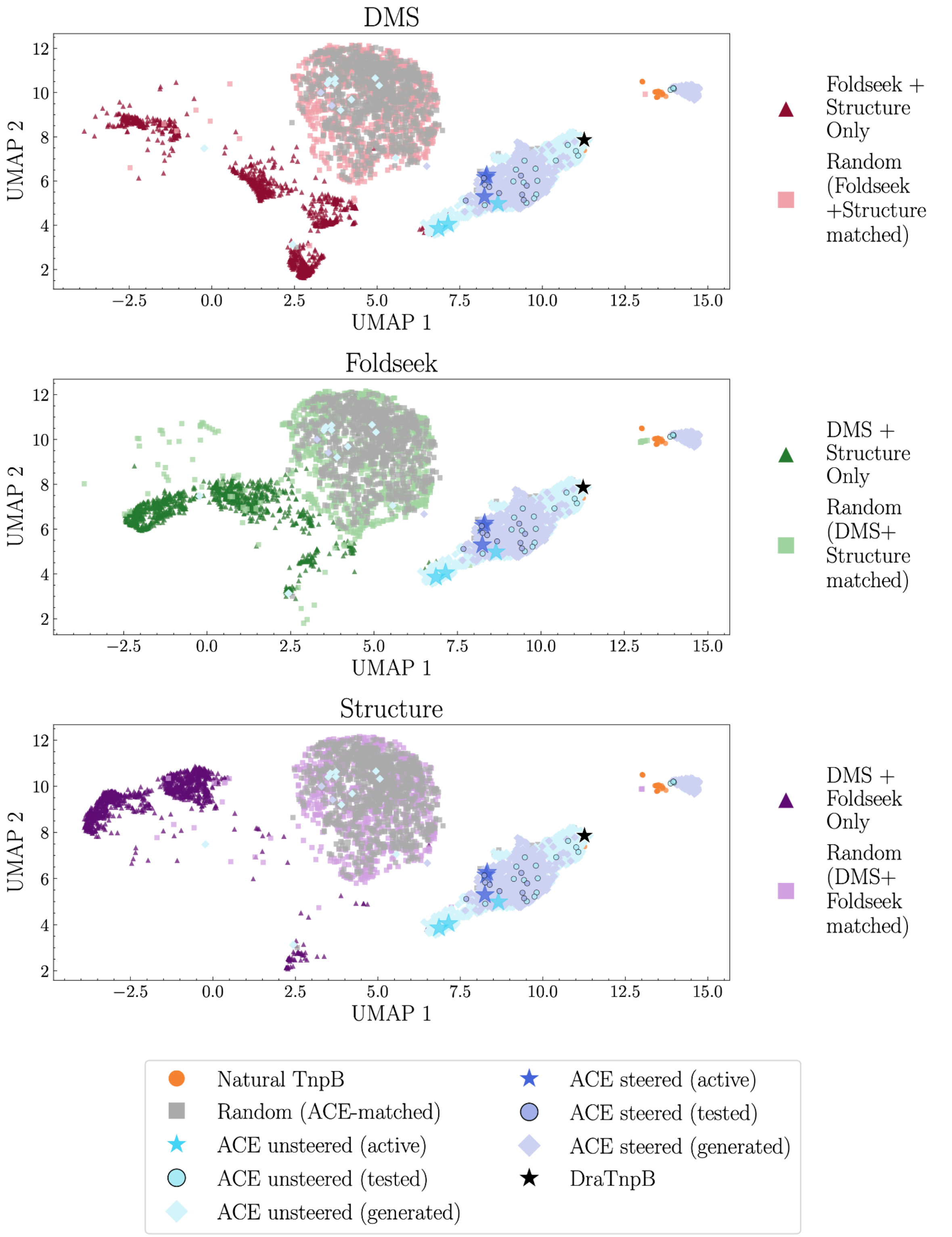


**Supplementary Figure 13. Contribution of evidence layers to ACE sequence phenotype.** For each evidence layer in turn (DMS [functional], Foldseek [evolutionary], Structure [structural]), the sites it nominated were removed from the input prompt, and the unsteered design pipeline generated editors using only the two remaining layers (dark-shaded triangles). Additionally, a random set of positions was set to wildtype in the input, with the number of fixed positions set equal to the number of sites captured by the remaining two evidence layers (light-shaded squares) or all three evidence layers (grey squares). DraTnpB, generated ACE constructs (using all three layers), and natural TnpB orthologs are also represented. Sequences were embedded with ESM2 650M. The full set of sequence embeddings were reduced to 50 dimensions via PCA and then projected onto a joint UMAP embedding. Each panel displays a filtered subset of the full UMAP embedding.

| Method | Effector / scaffold | Generative approach | Designs generated | Experimentally tested | Active | Hit rate (per design tested) | Hits per $10^6$ designs generated | Validation assay | Identity of active designs |
|---|---|---|---|---|---|---|---|---|---|
| **GenACE (this study)** | **Compact TnpB (ISDra2, 408 aa)** | **ESM-3 constrained masked infilling (functional + structural + evolutionary) ± Feynman–Kac steering** | **36,000** | **44** | **6** | **13.6%** | **167** | **Editing in human cells (HEK293T; GFP-ON reporter + endogenous loci)** | **57–61% to ISDra2 TnpB** |
| — steered | | with Feynman–Kac steering | 4,500 | 23 | 3 | 13.0% | 667 | | 57–58% |
| — unsteered | | without steering | 31,500 | 21 | 3 | 14.3% | 95 | | 60–61% |
| SynTnpB Skopintsev et al. | Compact TnpB (ISDra2) — same scaffold | ESM-IF1 inverse folding + evolution-conditioned masking + consensus averaging | not reported | 1,980 (bacterial) 9 (mammalian) | 466 (bacterial) 9 (mammalian) | 24% bacterial 9/9 mammalian | n.d. | Bacterial ccdB selection, then editing in HEK293T and Arabidopsis | 77–91% to ISDra2 TnpB |
| OpenCRISPR-1 Ruffolo et al. | Cas9 (SpCas9 family, ~1,380 aa) | ProGen2 fine-tuned on CRISPR-Cas Atlas + filters + language-model likelihood ranking | 350,000 | 209 | 131 any indel 48 clear indel | 62.7% any 23.0% clear | 137 | Editing in human cells (HEK293T; indels at 3 target sites) | 71.7% to SpCas9 |
| Evo — Cas9 Nguyen et al. | Cas9 locus (de novo, with sgRNA) | Evo genomic language model, autoregressive whole-locus generation | 2,000,000 | 11 | 1 | 9.1% | 0.5 | In vitro cleavage (no cellular editing) | 73.1% to SpCas9 |
| Evo — IS605 Nguyen et al. | TnpB transposon (IS200/IS605) | Evo genomic language model, autoregressive whole-locus generation | not reported | 24 | 3 | 12.5% | n.d. | In vitro excision / insertion (no cellular editing) | not reported |

**Supplementary Figure 14. Comparison of generative genome editor design methods.** Summary of various recent approaches in using screening and computational methods for novel editor generation. Note that we can only provide a lower bound for the number of generated / tested constructs for other studies based on what is included in the respective reports.

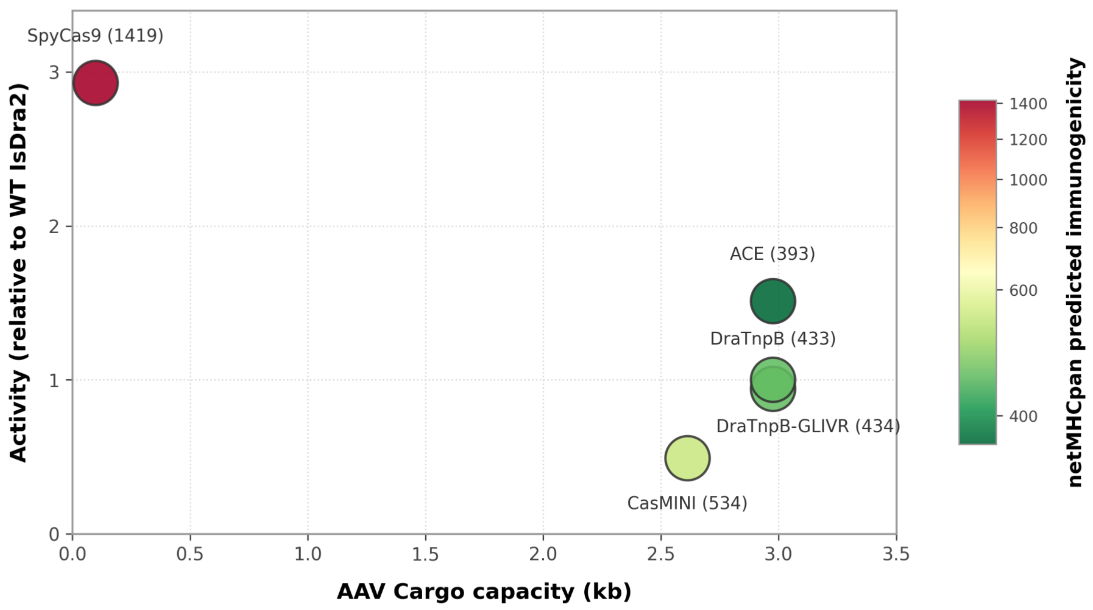


**Supplementary Figure 15. Comparison of gene editors across activity, cargo carrying capacity, and predicted immunogenicity.** For each effector, editing activity relative to DraTnpB averaged across 4 endogenous loci, is plotted against deliverability, as defined by remaining cargo capacity in AAV (kb). Bubble color is mapped to the sum of the number of predicted strong binders for MHC class 1 and 2, with exact values annotated in parentheses beside each label.
